# Implementation and System Identification of a Phosphorylation-Based Insulator in a Cell-Free Transcription-Translation System

**DOI:** 10.1101/122606

**Authors:** Shaobin Guo, Enoch Yeung, Richard M. Murray

## Abstract

An outstanding challenge in the design of synthetic biocircuits is the development of a robust and efficient strategy for interconnecting functional modules. Recent work demonstrated that a phosphorylation-based insulator (PBI) implementing a dual strategy of high gain and strong negative feedback can be used as a device to attenuate retroactivity. This paper describes the implementation of such a biological circuit in a cell-free transcription-translation system and the structural identifiability of the PBI in the system. We first show that the retroactivity also exists in the cell-free system by testing a simple negative regulation circuit. Then we demonstrate that the PBI circuit helps attenuate the retroactivity significantly compared to the control. We consider a complex model that provides an intricate description of all chemical reactions and leveraging specific physiologically plausible assumptions. We derive a rigorous simplified model that captures the output dynamics of the PBI. We perform standard system identification analysis and determine that the model is globally identifiable with respect to three critical parameters. These three parameters are identifiable under specific experimental conditions and we perform these experiments to estimate the parameters. Our experimental results suggest that the functional form of our simplified model is sufficient to describe the reporter dynamics and enable parameter estimation. In general, this research illustrates the utility of the cell-free expression system as an alternate platform for biocircuit implementation and system identification and it can provide helpful insights into future biological circuit designs.

## Introduction

The successful design and implementation of the inaugural biocircuits, such as the genetic toggle switch and the repressilator, have demonstrated the possibility of modularity in synthetic biological circuits [1, 2]. The recognition of functional modules makes building large and complicated biological circuits possible. Basic modules can be studied and tested in isolation and then can be connected with other modules to perform certain functions. However, the modularity of biological circuits can change when interconnections are made. This effect is called retroactivity and is a fundamental issue in systems engineering [3, 4]. This means that when a downstream system is connected to another system, the downstream system will affect the behavior of upstream component. As a result, the signal generated by the upstream component may not be effectively transferred to other components.

Retroactivity can be divided into two types based on which signal it affects, the retroactivity to the input and the retroactivity to the output. Based on previous theoretical studies, an operational-amplifier-like orthogonal biomolecular device could help attenuate retroactivity [3]. An electronic operational amplifier absorbs little current from upstream; as a result, there is almost no voltage drop to upstream output. At the same time, the retroactivity to the output is attenuated because of a large amplification gain and an equally large negative feedback loop. Based on these ideas and previous work [5], we tested an insulator design using nitrogen regulation proteins [6] in a cell-free transcription-translation (TX-TL) system.

The TX-TL system developed in [7, 8] is an attractive candidate platform for such rapid prototyping. The system facilitates DNA-based expression on plasmids and linear DNA, and since linear and plasmid DNA can be synthesized and expressed in the TX-TL system in a single day’s time [9], the time required to iterate over designs is considerably reduced.

Another powerful aspect of the TX-TL system is the ability to directly modulate the concentration of different pieces of DNA encoding different biocircuit components. The ability to rapidly synthesize and test the effect of different promoter sites, ribosome binding sites and other components, and simultaneously vary the DNA encoding these parts, permits a degree of freedom typically absent in cell-based assays. In this setting, iterating of prototypes can be assisted by predictive modeling of biocircuit dynamics. It is the ability to control DNA concentrations and rapidly vary structural properties of the biocircuit that allow us to address the problem of parameterizing a predictive model.

Cell-free systems have long been used to characterize fundamental parameters in biological systems [10]. In a synthetic biology context, especially for the phosphorylation-based insulator circuit, it is unclear what parametric information can be extracted from a series of systematic tests in an *in vitro* system, specifically the TX-TL system. With additional degrees of freedom in the experimental conditions, the TX-TL system may be able to provide insight into model parameters that *in vivo* studies could not. Moreover, it is unclear what systematic tests should be carried out in order to retrieve this information. This paper investigates these issues using the phosphorylation-based insulator as a case study.

In general, a parametric model is globally structurally identifiable only under certain mathematical conditions [11]. These conditions are valid as long as the control variables enter the dynamical system as a multiplicative perturbation. However, as we will see with the phosphorylation-based insulator, even if the model retains this structure the model may not be globally identifiable because of the large number of parameters it contains, despite having only a couple output variables. As is often the case, a first principles model may be physically representative of the intricate reactions happening in the system, but carry a complexity that far exceeds the information present in the data. Thus, simplified models that are reflective of the low-dimensional output data, while also retaining the (controllable) experimental variables in the TX-TL system, are desirable.

In this work, we successfully implement the PBI circuit in TX-TL and further propose a complex model based on the fundamental processes of transcription, translation, and phosphorylation. The model is unwieldy to analyze so we rigorously derive a simplified model based on a series of physically realistic assumptions, show that it is globally identifiable with respect to the data, and perform a series of experimental perturbation tests to back out the simplified model parameters.

The main contributions of this work are: 1) we demonstrate that the TX-TL system can be used to prototype relatively complicated synthetic biocircuits, such as the PBI circuit that involves not only transcriptions, translations and protein-DNA interactions but also post-translated interactions like phosphorylation and dephosphorylation; 2) we show that by utilizing the TX-TL system that has extra degrees of freedom compared to cell-based systems, we can systematically identify the parameters of our mathematical models using actual experimental data, which subsequently guide us to achieve more efficient circuit prototyping and better future circuit designs [12, 13].

## Demonstration of Retroactivity in TX-TL

Firstly, we wanted to demonstrate retroactivity in the TX-TL system. The example we used is a simple negative regulation circuit, in which constitutively expressed TetR proteins repress the transcription of downstream components pTet-GFP and pTet-RFP DNA unless an inducer aTc is added (Figure 1A). Here, we considered pTet-GFP as the reporter and pTet-RFP as the load. When there is no inducer present, the reporter will remain off because of the repression by TetR. However, if we added significant amount of load into the system, the load sequesters the TetR proteins from pTet-GFP, resulting in the activation of GFP transcription (Figure 1B). This is a result of retroactivity, in which downstream components affect the behavior of the upstream system output. We next tested this effect in the presence of different inducer concentrations (Figure 1C). At low aTc concentrations (less than 0.5 μg/mL), as load concentration increased, GFP expression increased because of retroactivity. However, if too much aTc was added, GFP expression actually decreased as load increased. This is because resources in TX-TL, such as ribosomes, RNA polymerase, are limited.

**Figure 1.**
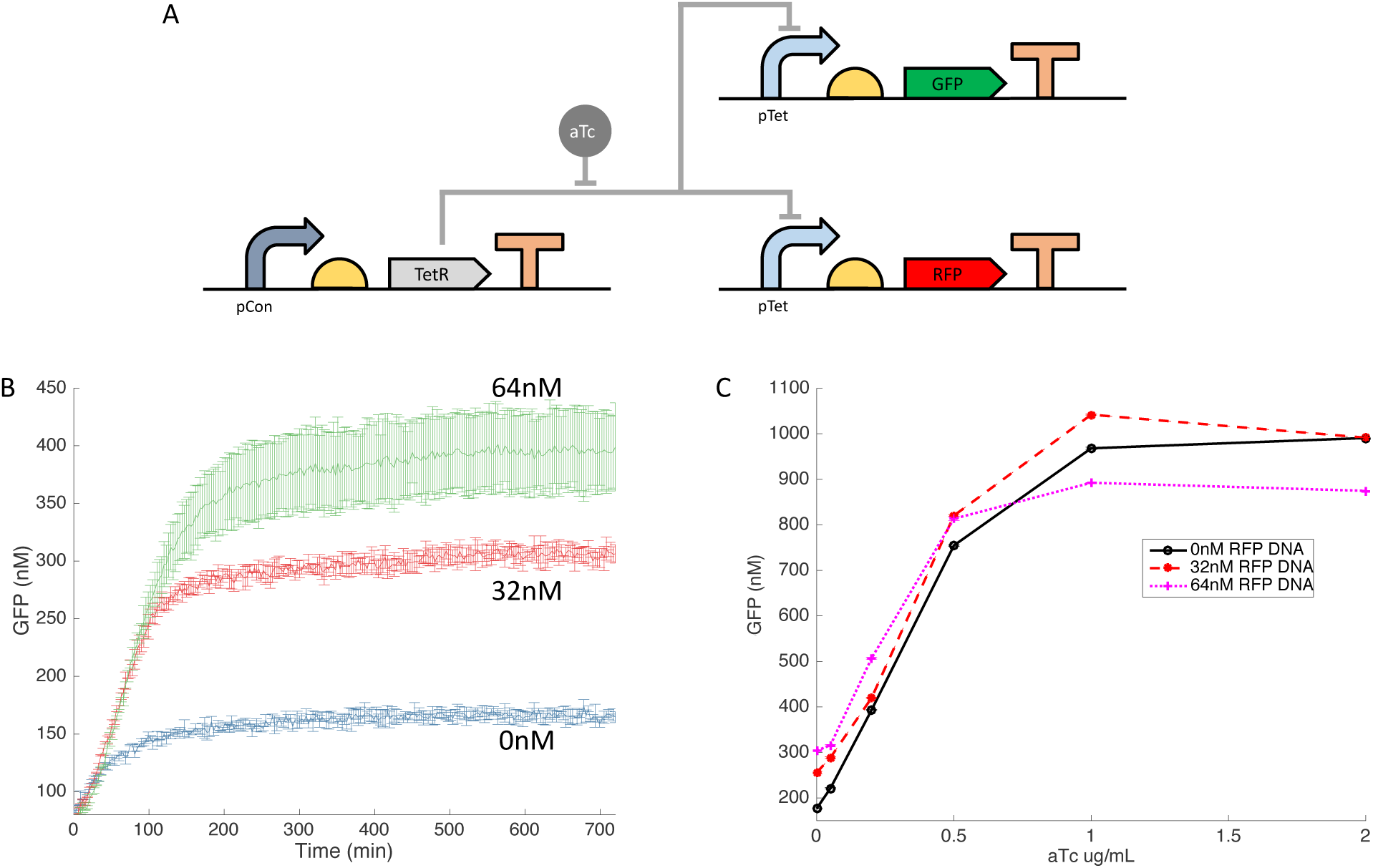
Demonstration of the retroactivity in the TX-TL system. **A**: Circuit diagram of a negative regulation circuit. **B**: Time traces of the GFP fluorescence in presence of different concentrations of RFP DNA. As RFP DNA concentrations increase, more GFP fluorescence can be detected as a result of the retroactivity. Error bars are standard deviations from 3 repeats. **C**: Titration ofTetR repressor aTc in presence of different concentrations of RFP DNA. X axis is the final concentrations of aTc in each sample and Y axis is the end point GFP fluorescence of the corresponding samples. Data were collected using a plate reader with settings for excitation/emission: 485 nm/525 nm.

This simple circuit demonstrates that there is retroactivity in biological circuits in the TX-TL system. To address this problem, we implement an insulator component to compensate for the retroactivity.

## Demonstration of the Insulation Capability of the PBI Circuit

Based on the insulator design in [5], we adapted a simpler form to implement in the TX-TL system (Figure 2A). The insulator design is based on a well-known two-component signal transduction system regulating the transcription of genes encoding metabolic enzymes and permeases in response to carbon and nitrogen status in *E. coli* and related bacteria [14]. There are two essential proteins in the system: NRII and NRI (NtrB-NtrC). NRI can be phosphorylated into NRI^P^ by NRII (kinase form). Only NRI^P^ is able to activate the σ54-dependent promoter glnA and trigger the transcription of downstream genes [15]. NRII is both a kinase and phosphatase, regulated by the PII signal transduction protein, which, on binding to NRII, inhibits the kinase activity of NRII and activates the NRII phosphatase activity [16]. NRII is known to form dimers and will autophosphorylate itself to become a kinase. Previous studies suggested that when NRII has a mutation of leucine to arginine at residue 16, it loses its phosphatase activity but shows normal autophosphorylation. In contrast, NRII with a H139N mutation is not able to transfer the phosphoryl group from its active site histidine to an aspartate side chain of NRI [14]. As a result, NRIIL16R only acts as a kinase and NRIIH139N only functions as a phosphatase.

**Figure 2.**
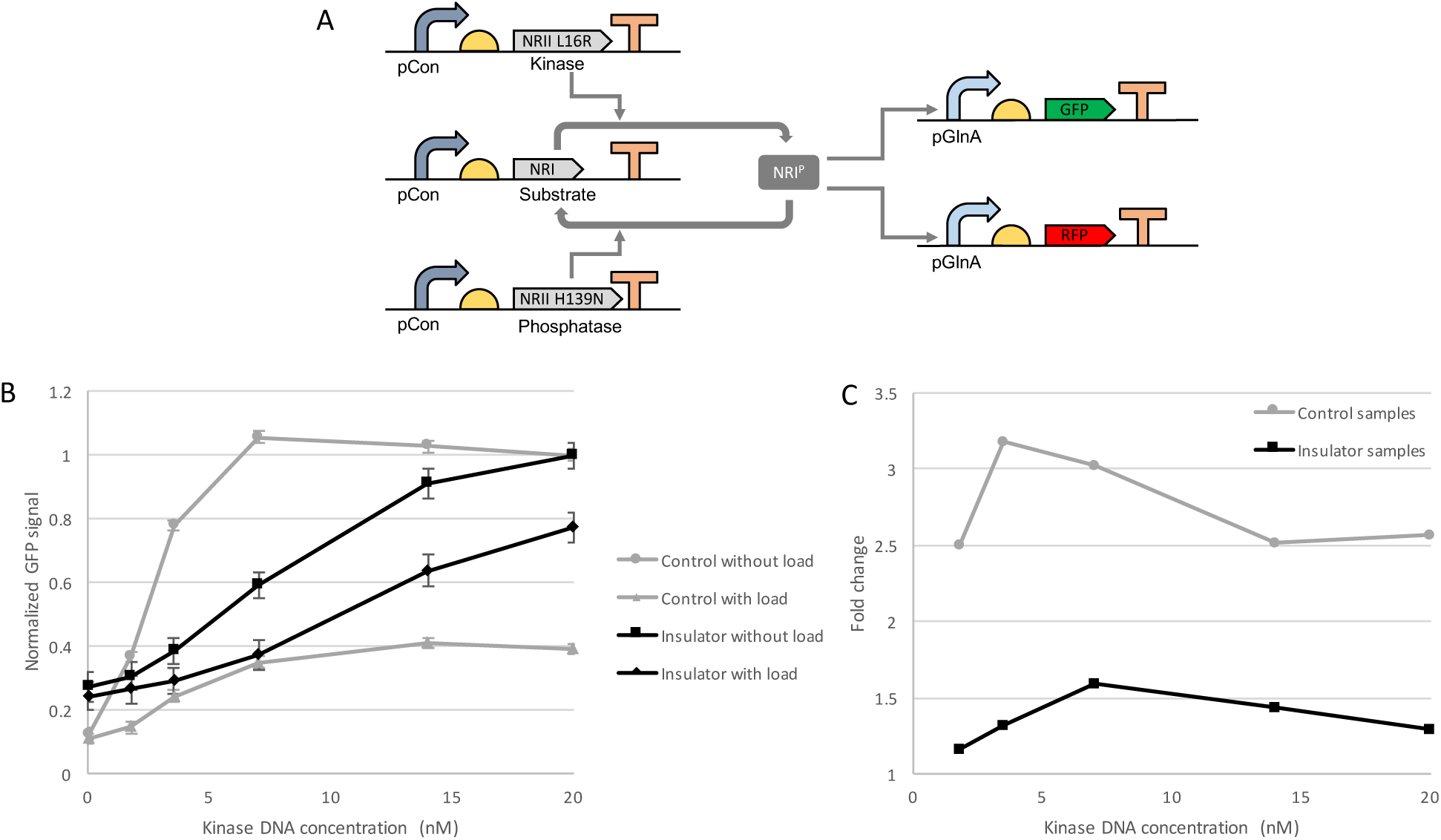
Implementation of the PBI circuit in the TX-TL system. **A**: Circuit diagram of the PBI circuit. pCon is a constitutive promoter. **B**: Transfer function curves for controls and insulators with or without load DNA. Raw GFP fluorescences were normalized using the highest GFP fluorescences from controls and insulators, respectively (highest GFP = 1). Compared to the control with load, which only had 40% signal left, the insulator with load was able to preserve 80% of the signal, significantly attenuating the retroactivity. **C**: Fold changes of the samples without load over the ones with load. The insulator samples have significantly smaller fold changes compared to those of the control samples.

In our circuit design, proteins NRI, NRIIL16R and NRIIH139N are all constitutively expressed. Reporter GFP is controlled by the σ54-dependent promoter glnA, which will be activated by phosphorylated NRI. We also confirmed that the phosphorylation-dephosphorylation loop worked in TX-TL (*Supplementary Figure S1*). By virtue of the fast timescale of phosphorylation and dephosphorylation loop, this circuit enjoys a large amplification gain and an equally large negative feedback mechanism as mentioned in the introduction, which makes it a promising insulation device.

To test the insulation capability of our insulator, we compared the behaviors of the insulator circuit with a control circuit that does not have a large amplification gain, neither a negative feedback loop. As mentioned in previous theoretical studies [3], the insulator circuit requires abundant substrate NRI to achieve high gain. So, we added 47.5 nM NRI linear DNA in the insulator circuit and only 5 nM in the control circuit. But to take resource limitations into account, we balanced the insulator and control by adding 42.5 nM of extra DNA (pTet-RFP) in the control circuit. Then we varied the amount of downstream glnA promoters by adding 0 nM (without-load) or 20 nM (with-load) pGlnA-RFP linear DNA, which would introduce retroactivity. We then titrated with different concentrations of NRIIL16R (kinase) linear DNA. After data were collected using a plate reader, the GFP relative fluorescence unit of the control and insulator circuits were normalized using their highest without-load fluorescence readings, respectively. As shown in Figure 2B, when the control circuit was added with load DNA, the GFP expression dropped significantly (about 60% at the highest kinase DNA concentration). In contrast, the insulator circuit only showed about 20% decrease in the GFP expression when the load was added at the highest kinase DNA concentration. Figure 2C simplifies the four curves into two curves by looking at the fold changes of without-load samples over with-load samples. As we can see, the insulator samples have much smaller fold change between without-load and with-load samples compared to the control samples, indicating the attenuation of the retroactivity by the PBI circuit. These results suggest that the insulator does help attenuate the retroactivity in biological circuits in TX-TL platform.

Besides, we also investigated the temperature sensitivity of this circuit in TX-TL. We performed the same experiments at 29°C, 33°C, 37°C, respectively (*Supplementary Figure S2*). At 29°C, the PBI circuit was able to limit retroactivity to about 20%; however, as temperature increased, the efficiency of insulation decreased to about 40% and 50%; while the control circuit had the same signal reduction among all three temperatures. The results suggested that the PBI circuit is sensitive to reaction temperature in TX-TL and its performance is affected by the temperature. Previous TX-TL characterization experiments suggested that relatively higher temperature would accelerate the molecular reactions involved in transcription and translation, resulting in faster GFP production rate [17]. However, because of the resource limitation in TX-TL, there could be more intensive competition on resources at higher temperature. As a result, the downstream load might have a better chance to sequester the NRI proteins away from the reporter at higher temperature, which would lead to less attenuation of the loading effect.

## Estimation of Constitutively Expressed Protein Concentrations

In this section, our goal is to derive a simplified model that can be uniquely parameterized from a set of characterization experiments in the bimolecular breadboard system [7, 8]. We base our model on the general phosphorylation-based insulator model posed in [3], but adapt our notation and augment input variables that are present in the biomolecular breadboard system. Because it is an *in vitro* system, the total DNA and inducer concentration in solution are adjustable experimental variables or variables that can be modeled as inputs. It is the freedom of these inputs that allows us to perform experiments and collect data that parameterizes the model.

We begin by introducing a chemical reaction model for the system:

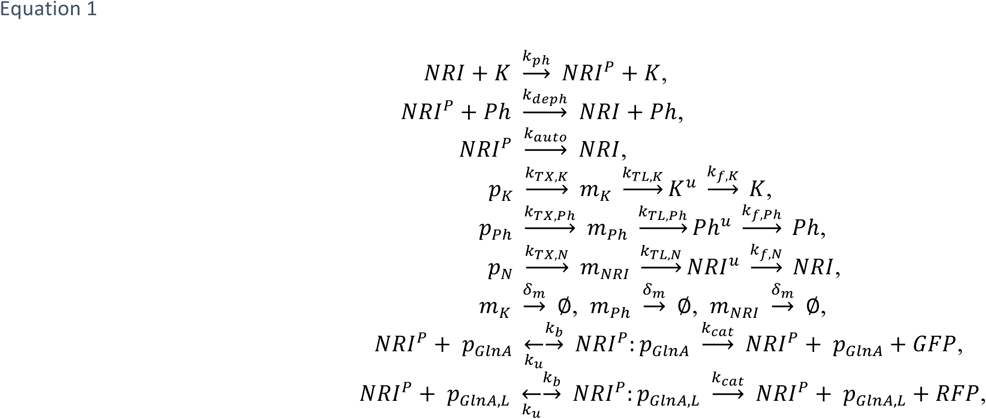
where *K, P, NRI, NRI*^*P*^ denote the kinase, phosphatase, unphosphorylated NRI and phosphorylated NRI protein, pGlnA is the GlnA promoter, pGlnA,L is the GlnA promoter encoding for other competing genes, *NRI*^*u*^ is the unfolded form of NRI protein, *K^u^* is the unfolded form of kinase, *Ph*^*u*^ is the unfolded form of phosphatase, and Ø represents a macro state of all degraded mRNA. We also use the notation *X*^*tot*^ when needed to denote the total amount of protein *X* where *X* = *K, Ph, NRI*. This notation will be convenient for our analysis in the sequel.

Since *p*_*x*_. represents a constitutive promoter for *X* = *K, Ph, NRl*, the total kinase, phosphatase, and NRI protein are produced constitutively. An assay with GFP shows that without additional proteases added into the bimolecular breadboard system, protein degradation is negligible [9]. Thus, we can approximate the total amount of NRI protein at a particular time *t* expressed under the pCon promoter using GFP expression expressed under the same promoter and ribosome binding site (RBS) as a proxy. This total amount of NRI, we will denote as *NRI*^*tot*^.

We also note that an alternative approach to estimate *NRI*^*tot*^ (*t*) is to assay the expression of a NRI-GFP fusion protein. However, this approach may significantly alter the phosphorylation dynamics of the NRI protein, since it acts as a substrate for the kinase. Therefore, we will express GFP separately on the pCon promoter with the same RBS and use it to estimate concentration from arbitrary units of fluorescence.

Because there are differences in the transcription and translation rates and folding of GFP and NRI, we do not expect the estimated concentration of GFP at time *t* will be identical to the concentration of the NRI protein at time *t*. We can account for these differences dynamically in a mass action model of NRI and GFP dynamics. If we consider NRI constitutive expression in a simple isolated system with no kinase or phosphatase activity *NRI*^*tot*^, e.g. with the chemical reaction system

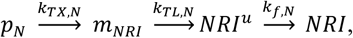
we see that 
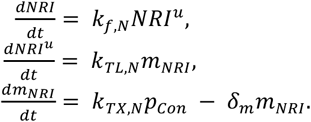

The total NRI protein at time *t* is ultimately a function of *m*_*NRI*_(*t*). Since the dynamics of *m*_*NRI*_ can be viewed as a scalar linear system with static step input pCon, we can solve analytically for *NRI*^*tot*^(*t*) to obtain:

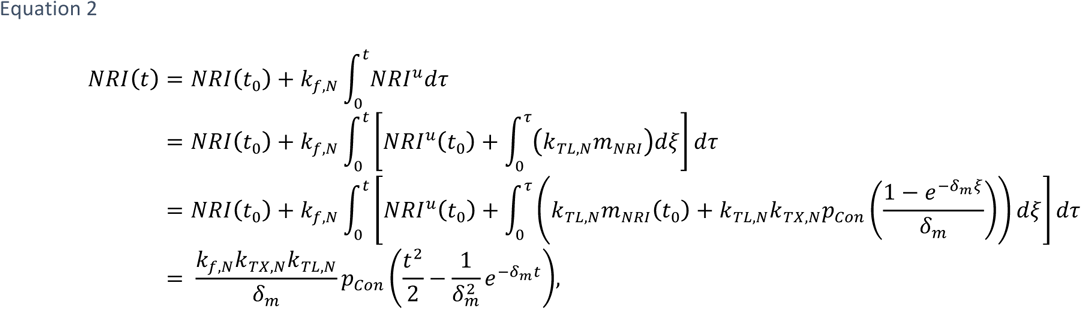
whenever *NRI*(*t*_0_) = *m*_*NRI*_(*t*_0_) = *NRI*^*u*^(*t*_0_) = 0. To reflect the experimental conditions of our system, we have assumed that the initial mRNA, unfolded and folded kinase, phosphatase and NRI concentrations are zero. Notice that in deriving this expression, we have made no assumption about time-scale separation. While such arguments are valid since the folding dynamics proceed at a much slower rate than the transcription and translation dynamics, they are unnecessary for estimating NRI at time *t*. Finally, it is worth noting that we assume the transcription and translation reactions proceed as first order reactions, which is valid as long as our DNA concentrations (typically in the nM range) are much less than the concentrations of RNA polymerases, ribosomes, chaperone proteins, etc. (typically in the μM range [8]).

It is worth noting that model for the mRNA species *m*_*NRI*_ is qualitatively consistent with our experimental studies of mSpinach expression in the transcription-translation system. To demonstrate this, we consider a model of the same functional form as *equation 2*, but with a constitutive promoter and coding sequence of the same length as the mSpinach transcript. This yields 
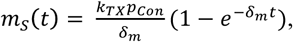
where 
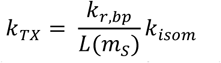
is estimated with *k*_*r,bp*_ = 60 *bp/s* (the approximate mean of a variety of media-dependent rates found in [18]), *L*(*m*_*s*_ = 98 *bp/nM* is the length of mSpinach aptamer without a tRNA scaffold [19], and *k*_*isom*_ = 6.3×10^−2^ *s*^−1^ is the forward rate of open complex formation from the closed complex.

From this, it is possible to estimate the rate of mRNA degradation,

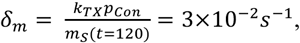
where *m*_*s*_(*t* = 120) is the expression of mSpinach at time *t* = 120 minutes and is an approximation of *m*_*s*_ steady state expression if the system were to continue to run indefinitely. The time point *t* = 120 minutes, is critical to consider for our biomolecular breadboard system. Previous results generated with the TX-TL system [9, 20] suggest that protein production rates typically maintain as a constant within 120 minutes of the reactions. After 120 minutes, we can see a decrease in mRNA concentration accordingly (Figure 3A) as well as slower protein production (Figure 1B). Thus, an empirical upper bound on time horizon for our model is approximately 120 minutes after the reaction is initiated.

**Figure 3.**
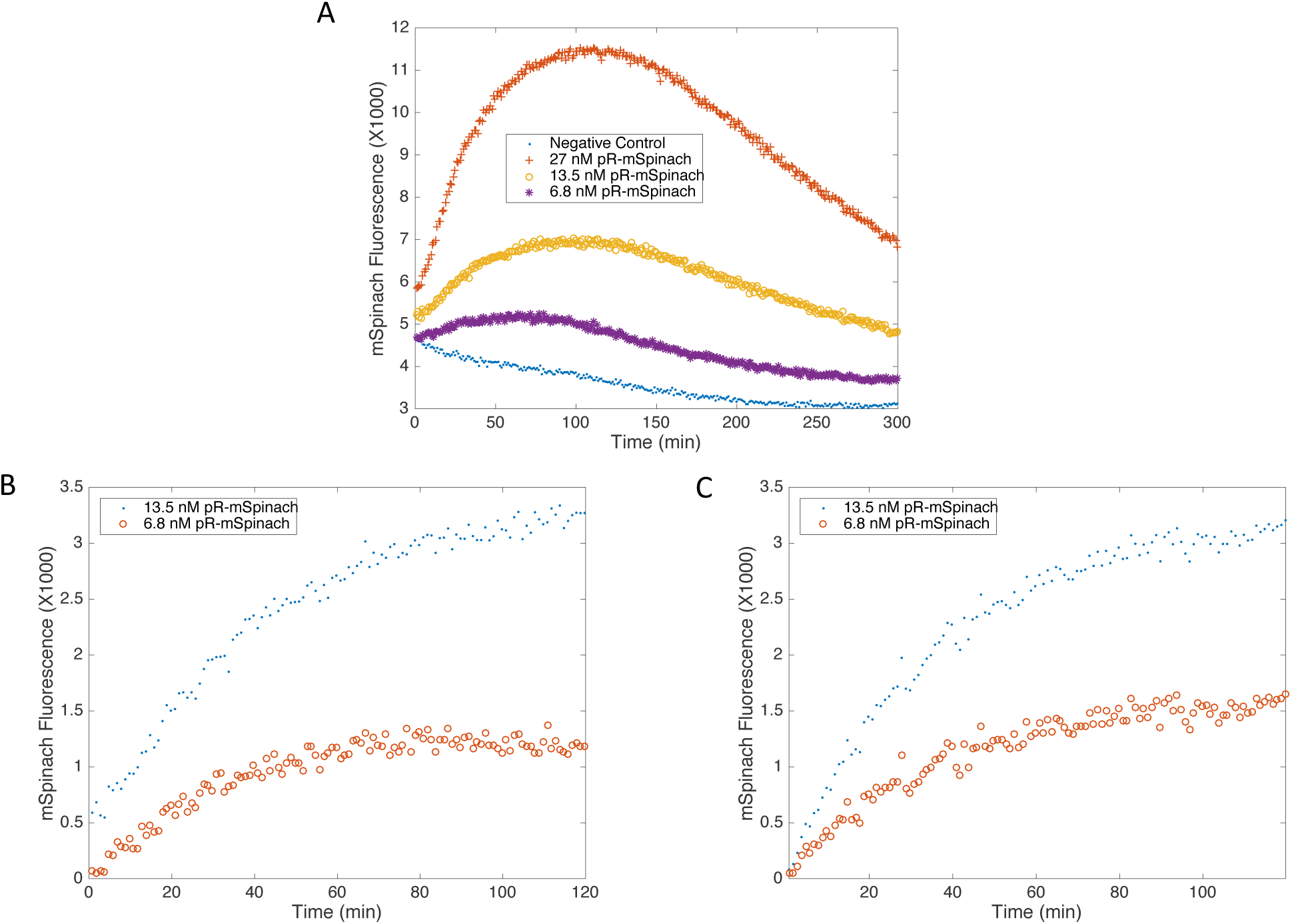
**A**: Data featuring mSpinach expression on linear DNA with 100 bp of protection. The transcriptional unit consists of an OR1-OR2-pR promoter, followed by the mSpinach (no scaffold) RNA aptamer coding sequence and the T500 terminator. Arbitrary fluorescence units of mSpinach expression is plotted against time. Subtracting the background, we see that mSpinach expression nearly doubles as DNA concentration doubles. Past *t* = 120 minutes, mSpinach expression decreases, presumably because linear DNA template has degraded or transcriptional resources are exhausted. Our time horizon of interest for the model will thus be in the interval of *t ∈* (0, 120). **B**: Data featuring mSpinach expression driven by the OR1-OR2-pR promoter at 13.5 nM and 6.8 nM concentration from the time interval of 0 to 120 minutes. mSpinach expression dynamics in the time horizon of interest feature a phase of steep linear growth and then saturation towards an asymptotic limit. **C**: A simulation of mSpinach expression, driven by a constitutive promoter at 6.8 nM and 13.5 nM DNA concentration. Notice that the model is able to capture the qualitative effects of mSpinach expression.

It is also important to mention that with the exception of the mRNA species *m*_*NRI*_ of NRI protein in our model, the species associated with NRI do not settle at a stable steady state. This aspect of our model is consistent with the behavior of biocircuit expression for an initial window of time in the biomolecular breadboard system. In this *in vitro* system, the auxiliary proteins NRI, K, Ph and even GFP (expressed by pGlnA) do not achieve a steady state in the traditional manner (due to detailed balance of production and a combination of degradation and dilution effects). Rather, they continue to increase in resources are exhausted. Thus, because we are interested in the dynamic behavior of the phosphorylation-based concentration until all transcriptional and translation insulator and drawing comparisons of its *in vitro* behavior to *in vivo* behavior, we will focus our subsequent modeling efforts in the time window *t ∈* (0,120) minutes where fuel, energy and other transcriptional and translational resources are still abundant. In doing so, we do not preclude the possibility of genes competing with each other for the finite resources available in the *in vitro* system. Our time frame of interest is thus when transcriptional and translational machinery is available and functional, but in finite supply (mimicking *in vivo* conditions).

Using the parameters that we have calculated, we plot the outcome of a simulation against expression data for mSpinach in Figure 3. The output of the simulation is simulated with additive white noise, replicating the measurement noise present in the plate reader (refer to the trajectory of the negative control). We use a biocircuit expressing mSpinach with the constitutive promoter (pOR1-OR2 from the λ regulatory operon). Notice that the functional form of *m*_*NRI*_(*t*) adequately describes the qualitative behavior of mRNA expression in the breadboard system until *t* = 120 minutes. The rate at which mSpinach saturates is determined by the *δ*_*m*_ parameter and its steady state value is given as *k*_*TXpCon*_/*δ*_*m*_. These experiments with the mSpinach RNA aptamer show that our model, while simple in its formulation, is sufficiently complex to describe transcriptional dynamics in the transcription-translation system for the first two hours. Thus, we will not attempt to model system expression when the transcription-translation system depletes it resources; at this point gene expression is strongly competitive, production and degradation rates are largely determined by the available ATP, rNTP, amino acids, etc. in the system.

We also know that the folding rates of the GFP protein are different from those of NRI protein. Thus, to estimate the ratio in folding rates, we use the K-fold protein folding simulation software developed in [21]. Based on previous studies [22], we use the following equations to estimate the mRNA transcription rate and the protein translation rate: 
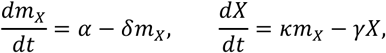
where *m*_*X*_ is the concentration of mRNA for protein *X* (*X = NRI, GFP*), *α* is the rate of production of the mRNA for protein *X, δ* the rate of degradation of the mRNA, *κ* is the rate of translation of mRNA and *γ* is the rate of degradation of protein *X*. The value of *α* increases with the strength of the promoter while the value of *κ* increases with the strength of the RBS [22]. Because we are using the same promoter and RBS for both NRI and GFP genes, they should share the same mRNA transcription rate and protein translation rate, despite their differences in gene length.

We express the rates of transcription, translation, and folding for NRI in terms of GFP rates of transcription, translation, and folding (respectively) as follows:

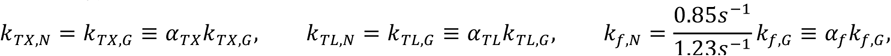
where “≡” denotes a definition of *α*_*j*_ Our model for GFP expression under the pCon promoter is similarly expressed as

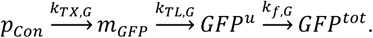

The model derived for *GFP*(*t*) follows an analogous derivation as the model for *NRI*^*tot*^(*t*). Thus, using *equation 2*, it is straightforward to show that *NRI*^*tot*^(*t*) concentration can be expressed as

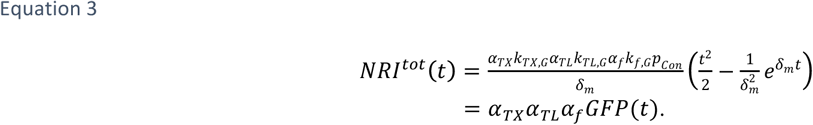

We see that by scaling the GFP concentration by the appropriate ratios at time *t*, we can obtain an estimate for *NRI*^*tot*^. The above formula holds as long as the concentration of pCon promoter expressing GFP is the same as the concentration of pCon promoter expressing NRI protein with the same RBS. Otherwise, a ratio to account for the scaling between the two should also be incorporated into the above relation.

To summarize, we have posed a basic model for constitutive expression of NRI protein; the model has a closed form analytical expression that allows estimation of total NRI protein as a function of time. Our model relies on a basic set of chemical reactions describing the processes of transcription and translation. To justify our model at the transcriptional level, we have performed an experimental assay using the mSpinach RNA aptamer to ascertain the dynamics of mRNA expression in the biomolecular breadboard system. Our simulations and experimental data appear to match for up to the first two hours of the experiment, based on parameters extracted from various references, suggesting that our model is accurate in a time horizon of interest. We thus restrict our attention to this time horizon, as it represents the horizon in which transcription and mRNA degradation proceed unperturbed. Further, evidence in [17] suggests that ribosomal activity proceeds unhindered in the first two hours.

We also observe that an analogous line of reasoning can be applied to estimating *Ph*^*tot*^ and *K*^*tot*^. We do not repeat the derivation here, as it only requires a change in notation. However, we emphasize that because of these observations, in the sequel we will refer to *Ph*^*tot*^(*t*), *K*^*tot*^(*t*), and *NRI*^*tot*^(*t*) as additional input variables (so long as we are modeling the appropriate time horizon). Additionally, it is the ratio of *Ph*^*tot*^ and *K*^*tot*^ that matter as a functional input in the system identification process and not the individual concentrations that matter. Further, it is by levering the inputs *Ph*^*tot*^/*K*^*tot*^, and *NRI*^*tot*^ we are able to identify the parameters of a simplified model uniquely.

## Derivation of A Simplified Model for the PBI

In this section, we derive a simplified model of the phosphorylation-based insulator using the chemical reaction system (*equation 1*). Examining the full chemical reaction system (*equation 1*), we obtain the following state space model from the law of mass action:

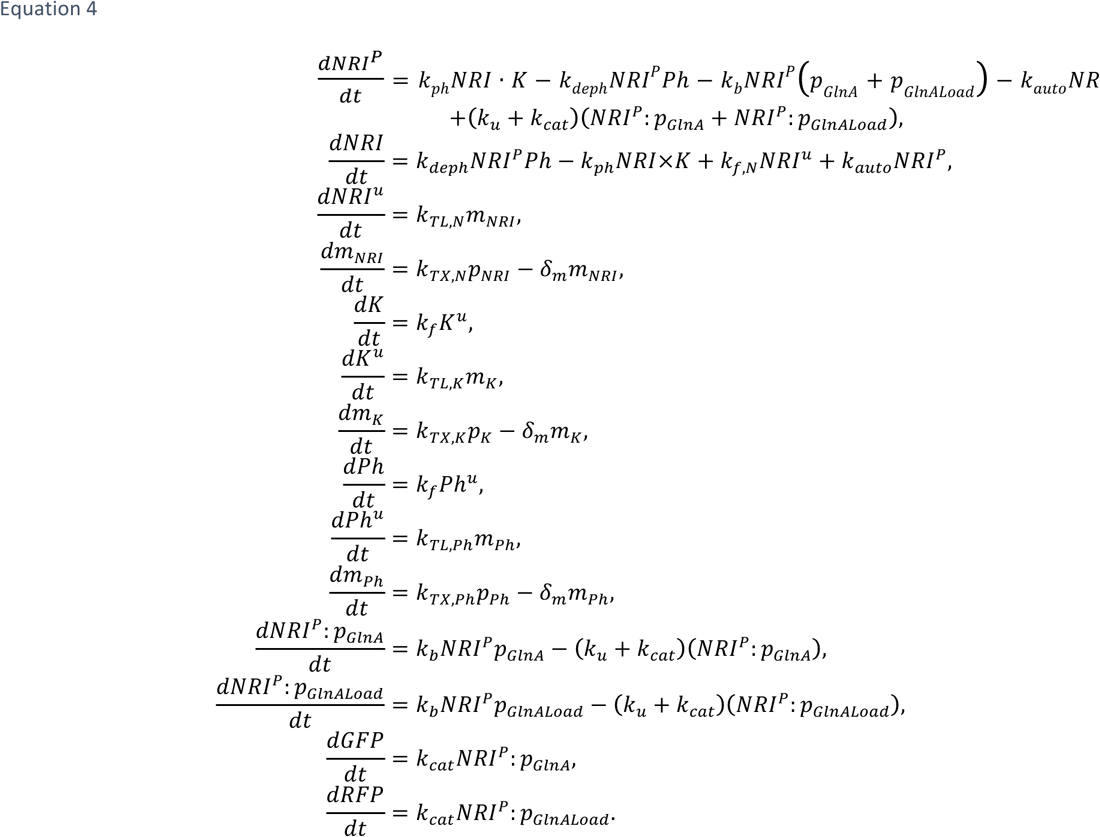

The dimension of the state-space model is 14 and because of the presence of bimolecular reactions, it is nonlinear in the state of the system. Thus, it is difficult to obtain a closed form expression for the solution to the system. However, we will systematically impose a series of modeling assumptions that are physiologically plausible, but which greatly reduce the complexity of the model.

First, notice that the total concentration of K, Ph and NRI, denoted as *K*^*tot*^, *Ph*^*tot*^ and *NRI*^*tot*^, depends only on the transcription and translation reactions. Thus, if we consider the transcription and translation dynamics of K, Ph and

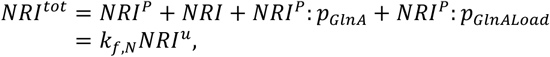
in isolation, we can use the results of the previous section to obtain a closed form expression for their total concentration as follows:

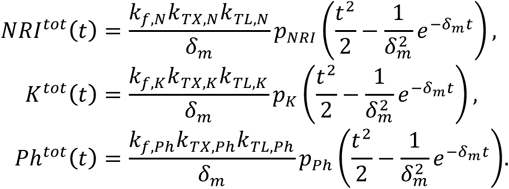

These total concentrations can be viewed as time varying parameters. If we had a way of quantifying the rate of transcription, translation, and folding of the individual proteins in the transcription-translation system, we could predictively estimate the trajectories of *NRI*^*tot*^, *K*^*tot*^ and *Ph*^*tot*^ over time. However, we do not have these parameters, and thus it is advantageous to employ the previous section’s approach. With similar arguments, we can argue that the total concentration of these proteins can be expressed as the total concentration of a reporter molecule multiplied by a scaling constant (see *equation 3*). Thus, using a separate assay to quantify constitutive expression of a reporter molecule under a given constitutive promoter (and a calibration curve to convert fluorescence to molar concentration), we can use the reporter molecule as a proxy for estimating the true molar concentration of NRI, kinase or phosphatase. Therefore, we can avoid the problem of estimating transcriptional, translational and folding rates of heterogeneous proteins while obtaining an estimate of the functional protein concentrations. Moreover, the result holds for all t in which RNA expression increases linearly (*t* ≤120).

We formalize this assumption as follows:

### Assumption 1

We suppose that for all *t ∈* [0,120], *K*^*tot*^(*t*), *Ph*^*tot*^(*t*), and *NRI*^*tot*^(*t*) are known parameters. This assumption thus allows us to eliminate the dynamics of folded kinase, unfolded kinase, folded NRI, unfolded NRI, folded phosphatase, unfolded phosphatase and all mRNA dynamics.

The remaining dynamics of the system are thus given as:

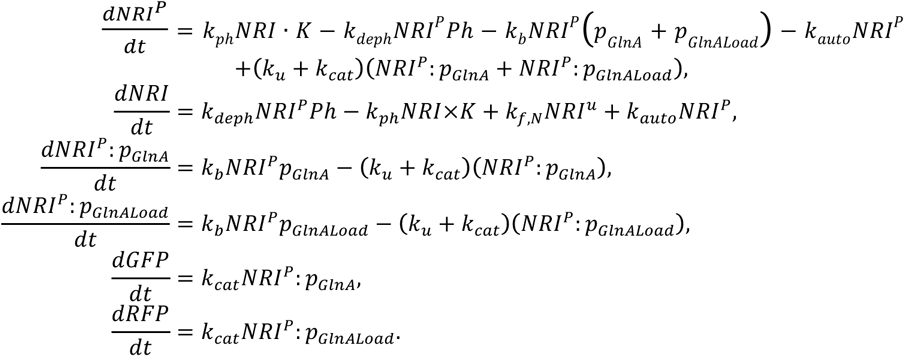

Next, we assume that the phosphorylation and dephosphorylation reactions occur at a much faster time scale then production of GFP or RFP and the binding (and unbinding) reactions of NRI^P^ to DNA to form (or disintegrate) activator-DNA complex. We justify the latter assumption through experimental observations that observe phosphorylation rates on the order of 10^6^ *min*^−1^. Transcription factor binding rates are less characterized but typically binding and unbinding rates of a transcription factor (e.g. LacI) are *O*(10^−1^) *min*^−1^ and *O*(10) *min*^−1^ respectively [23].

We formalize these assumptions as follows:

### Assumption 2

We suppose that *k*_*ph*_, *k*_*deph*_ ≫ *k*_*u*_, *k*_*cat*_, *k*_*b*_, *k*_*auto*_.

Next, we suppose that the amount of DNA bound NRI^P^ is smaller than the amount of free NRI^P^ and unphosphorylated NRI and that total NRI can be approximated as the sum of unbound NRI^P^ and NRI. Put another way, we assume that the molar concentration of unbound NRI protein is substantially larger than the molar concentration of DNA-bound NRI protein. This will certainly be the case since the pGlnA and pGlnAload DNA concentration will be in the nM range while the protein concentration of NRI will be in the μM range (refer to the arguments in the previous section and Figure 3). From the above reactions and assumptions, we then can write the dynamics of NRI^P^ using the approximate conservation law

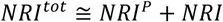
as follows:

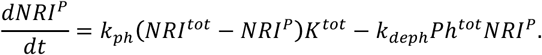

Since phosphorylation and dephosphorylation occurs at a much faster rate than GFP and RFP production (our ultimate time-scale of interest) and reasonably faster than the binding dynamics of the NRI^P^ transcriptional activator, we can solve the fast dynamics of NRI^P^ to obtain an analytical expression for the equilibrium point 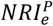. At steady state, we have

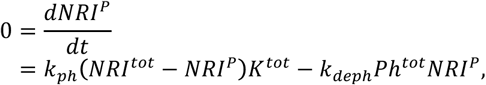
which implies

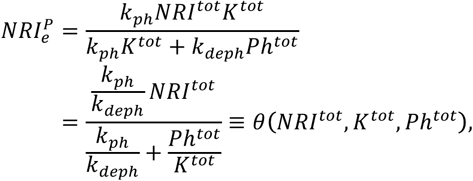
where “≡” denotes the definition of the function *θ*(*NRI*^*tot*^, *K*^*tot*^, *Ph*^*tot*^).

In this assumption, we also assume that *k*_*auto*_ is negligible compared to other rates. This is a reasonable assumption since spontaneous dephosphorylation proceeds at a slow rate — the ΔG of spontaneous dephosphorylation is very large [24].

The final assumption we leverage is that the rates of GFP and RPF production, relative to the binding dynamics of *NRI*^P^ are much slower. Specifically, we suppose that:

### Assumption 3

*k*_*b*_, *k*_*u*_ ≫ *k*_*cat*_.

This assumption can be justified, since the production of a folded protein such as GFP takes at least ten to fifteen minutes [25] while the binding and unbinding rates are typically on the order of hundredths of seconds and seconds, respectively [23]. Thus, we can solve for the steady state of the DNA-activator complexes NRI^P^:pGlnA and NRI^P^: pGlnALoad. The result is analogous to the classical Michaelis-Menten model, with 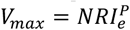 and *K*_*m*_ = (*k*_*u*_ + *k*_*cat*_)/_*kb*_. We omit the derivation, as it follows the standard derivation for a two-substrate one-enzyme model:

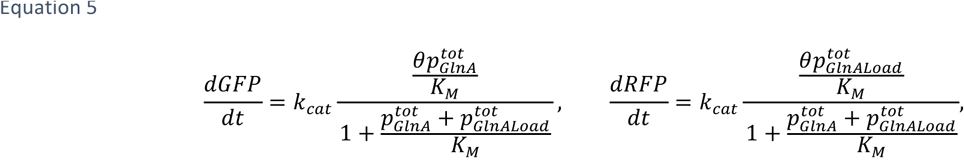
where *θ* denotes *θ*(*NRI*^*tot*^, *K*^*tot*^, *Ph*^*tot*^). This completes the derivation of our simplified model. In the next section, we will explore the analyze the structure of the model, determine which of the parameters are globally identifiable, and under what circumstances identifiability holds.

## System Identification of the Simplified PBI Model

### A. Theoretical Analysis

In the derivation of our model we have made a point to retain the experimental parameters *NRI*^*tot*^, *K*^*tot*^, *Ph*^*tot*^, 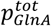 and 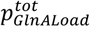. These parameters can be viewed as experimentally controllable, in that we can directly control the DNA concentration of promoters 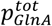 and 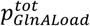. Additionally, by adjusting the underlying constitutive promoters driving the expression of NRI, K, and Ph we can effectively tune the quantities *NRI*^*tot*^, *K*^*tot*^ and *Ph*^*tot*^. We note this type of control over the concentration of DNA as well as total protein concentrations is not typically achievable in vivo, unless inducers are employed or different replication origins are cloned into a plasmid (which introduces variability in copy number from cell-to-cell). However, this advantage in the biomolecular breadboard is precisely the capability required to explore the problem of parameter estimation and determine if our simplified model is globally identifiable.

Since our calibration curves allow us to estimate GFP concentration from arbitrary fluorescence units, we will focus our attention on the GFP dynamics. Furthermore, notice that the forms of dynamics of both reporter molecules are identical. Thus, it suffices to analyze the identifiability of parameters with respect to the output dynamics of the GFP reporter molecule, since it will yield the same result as studying identifiability with respect to RFP output dynamics. Recalling our assumptions from the previous section, we will also make a point to study the behavior of the system within the time horizon of interest captured by our model, *t ∈* [0, *τ*_*max*_) where *τ*_*max*_ is the initiation of the resource depletion phase in our transcription-translation system.

Our goal is to determine whether this model is globally structurally identifiable with respect to the parameters *K*_*m*_, *k*_*cat*_, *k*_*ph*_, and *k*_*deph*_, given the inputs *NRI*^*tot*^, *K*^*tot*^, *Ph*^*tot*^, 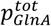 and 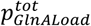. Notice that the inputs do not enter the dynamics of the system in a linear fashion. Indeed, the simplified system (*equation 5*) is of the form: *ẋ* = *f*(*U*, Θ), where *f* is nonlinear with respect to *U* and Θ, in which 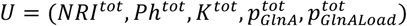 and Θ = (*K*_*M*_, *K*_*cat*_, *k*_*ph*_, *k*_*deph*_).

Furthermore,

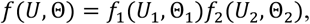
where 
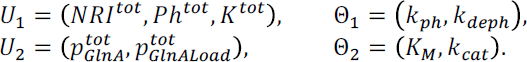

Notice that 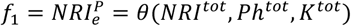 takes the form of a Hill function with Ph^tot^/K^tot^ as its substrate and

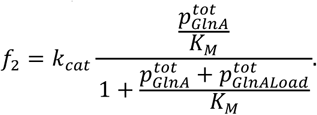

This multiplicative decomposition provides a key insight: our system dynamics is the product of two Hill functions with distinct inputs for each Hill function. This suggests that from a system identification standpoint, we can attempt a series of experiments that perturb one of the Hill functions while holding the other constant and vice versa to tease out the parameters for each.

To obtain insight into the what parameters in the Hill functions are identifiable, we invert the system dynamics to obtain

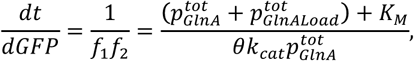
which indicates that the parameter identification is a linear regression problem and after some arrangement, we obtain that 
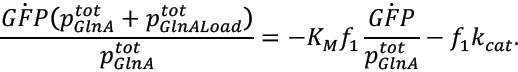

Thus, when the experimental input 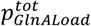 is set to 0 nM, we obtain a linear regression problem in estimating slope *K*_*M*_*f*_1_ and intercept *k*_*cat*_*f*_1_. Further, if we enforce that *Ph*^*tot*^ = 0, then *f*_1_ reduces to *NRI*^*tot*^, a known input value that completes the decomposition. Thus, by enforcing these two input constraints, we obtain a linear regression problem that effectively estimates *k*_*cat*_ and *K*_*M*_. By varying the total DNA concentration 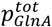 we can thus vary the rate of change of GFP, *dGFP/dt*, and obtain data to optimize *k*_*cat*_ and *K*_*M*_. Once *k*_*cat*_ and *K*_*M*_ are estimated, we can then use a similar line of arguments to back out an estimate for the ratio *k*_*ph*_/*k*^*deph*^.

In particular, we consider a nominal operating concentration of 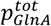, 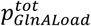 and write *γ* = (*U*_2_,Θ_2_)/*f*_2_ and *k*_*r*_ = *k*_*ph*_/*k*_*deph*_, then taking the reciprocal of dGFP/dt we obtain

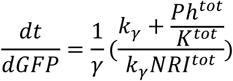
and define 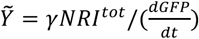 we see that

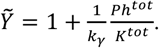

Therefore, by transforming the problem into the reciprocal space, we see that *k*_r_ = *k*_*ph*_/*k*_*deph*_ is a uniquely identifiable parameter. That is, the problem of estimating *k*_*γ*_ can be expressed as a linear regression problem with *k*_*γ*_ as the reciprocal of the slope and an intercept of unity. The fact that we can write the parameter estimation problem for (*k*_*cat*_, *K*_*M*_, *k*_*ph*_/*k*_*deph*_) as a solution to a system of linear equations thus shows that the model is globally structurally identifiable with respect to (*k*_*cat*_, *K*_*m*_, *k*_*ph*_/*k*_*deph*_).

In summary, we have derived a simplified model for the phosphorylation-based insulator and shown it is globally identifiable with respect to the output trajectory of GFP. We have shown that in the theoretical scenario where a continuous trajectory of GFP can be obtained to estimate its derivative *dGFP/dt*, the parameters *k*_*cat*_, *K*_*M*_ and *k*_*r*_ = *k*_*ph*_/*k*_*deph*_ can be estimated. These parameters are only estimated through a series of carefully designed experiments in which specific TX-TL controllable experimental variables are tuned. In the next section, we discuss the results of these experiments and numerical estimation of this data from time-series data.

### B. Experimental Analysis: Systematic Perturbations of the Phosphorylation-Based Insulator for System Identification

To identify the parameters *k*_*cat*_, *K*_*M*_ and *k*_*γ*_, we needed to perturb the phosphorylation-based insulator with the experimental variables designated in our model. In particular, we first needed to perturb the amount of pGlnA promoter producing GFP in the absence of phosphatase *Ph*^*tot*^ or NRIIH139N protein. Varying the amount of pGlnA promoter in the system in the absence of phosphatase would enable the estimation of *k*_*cat*_ and *K*_*M*_.

Intuitively, *k*_*cat*_ and *K*_*M*_ characterize the enzyme-substrate relationship that the activator protein NRI^P^ has with the pGlnA promoter — coincidentally, to reveal these parameters we need to eliminate any negative feedback imposed on the activator protein by NRIIH139N phosphatase and vary the substrate concentration 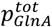 to reveal the kinetic parameters.

Accordingly, we ran a set of TX-TL reactions in which the DNA concentration of pCon promoter driving NRIIH139N expression was 0 nM. We varied the concentration of pGlnA promoter from 0 to 57 nM, expressed on plasmid. From the time series data, we extracted the first thirty minutes of expression dynamics — this time horizon constituted the time frame when amino acids, CoA, NADH, ATP, etc. were far away from the stage of complete depletion in the TX-TL system. In this time horizon of interest, the expression of GFP is linear with respect to time; therefore, the derivative of GFP is constant and can be fitted using the slope of a linear regression. The results of our linear regression are plotted against the time series data of the experiment in *Supplementary Figure S3*. The estimates of the *dGFP/dt* in the time horizon of interest at varying concentrations of pGlnA were used to fit the Hill function parameters *k*_*cat*_ and *K*_*M*_:

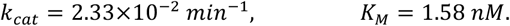

We emphasize that the key to estimating *k*_*cat*_ and *K*_*M*_ is the additional freedom afforded by a control input 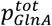 in perturbing the system.

The final parameter to estimate was *k*_*r*_ = *k*_*ph*_/*k*_*deph*_. In order to estimate *k*_*γ*_, we needed to fix the pGlnA concentrations, i.e. the concentrations driving expression in the downstream module and, and perturb the phosphorylation-based insulator. Specifically, we varied the ratio of kinase (NRIIL16R) to phosphatase (NRIIH139N) in the system by varying the ratio of DNA concentrations for the promoters driving their expression. Doing this, we obtained a series of time-lapse curves of GFP expression over a range of *Ph*^*tot*^/*K*^*tot*^ values. Again, we extracted estimates for *dGFP/dt* using a linear regression over the first thirty minutes of gene expression. The resulting estimates for dGFP/dt were then plotted against varying *Ph*^*tot*^/*K*^*tot*^ to fit a Hill function, see Figure 4. Using standard linear regression techniques, we then obtained the following estimate:

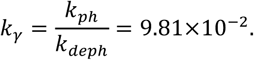

**Figure 4.**
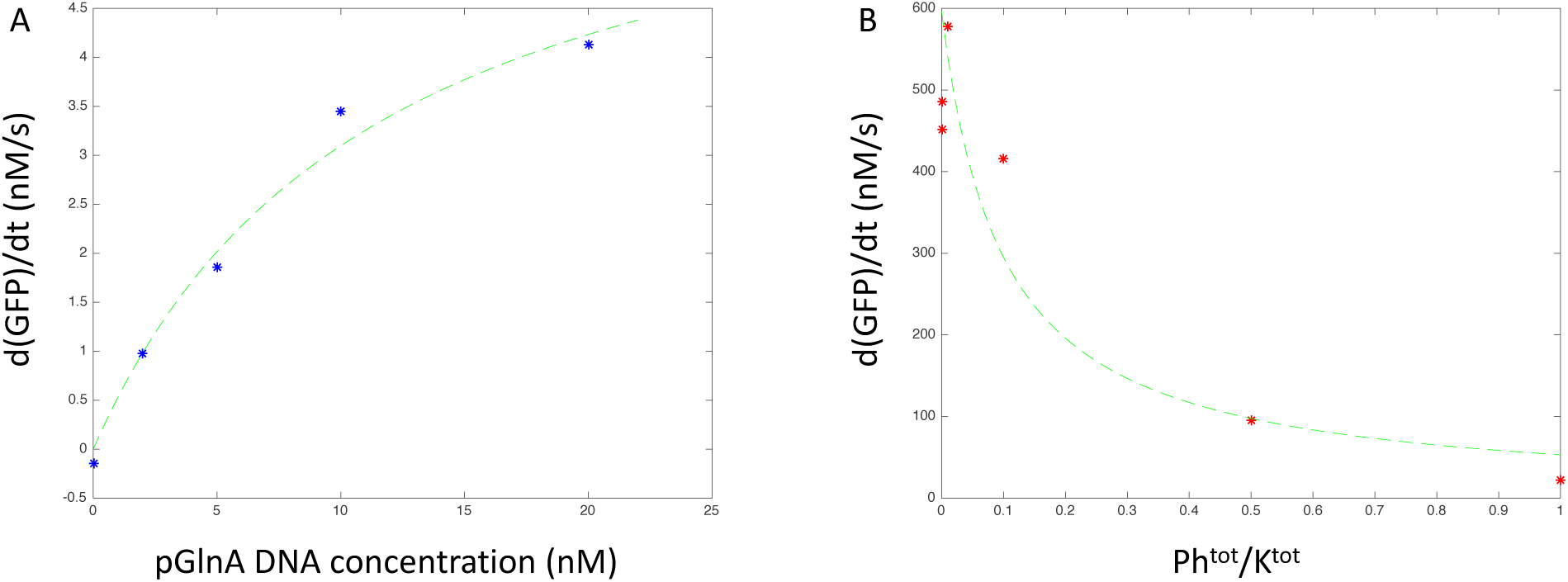
**A**: A plot of the resulting Hill function *dGFP/dt* against varying pGlnA. We see the curve follows the form of a Michaelis Menten function which is consistent with our model. **B**: A plot of the resulting Hill function *dGFP/dt* against varying *Ph*^*tot*^/*K*^*tot*^. Again, the empirical data (starred) matches the functional form of our model.

The ratio *k*_*ph*_/*k*_*deph*_ characterizes the balance of power between phosphorylation and dephosphorylation reactions — although we are unable to infer the individual parameters *k*_*ph*_ and *k*_*deph*_ we are able to conclude that dephosphorylation occurs at roughly an order of magnitude faster than phosphorylation (all other variables equal). Notice this parameter characterizes the intrinsic chemical reaction rates, rather than the flux or mass action rates that are dependent on kinase and phosphatase concentrations. Thus, to tune the phosphorylation-based insulator we can vary the amount of kinase and phosphatase concentrations, bearing in mind that phosphorylation is slightly slower than dephosphorylation in the TX-TL system.

Further, it is consistent with our intuition that only the ratio of *k*_*ph*_ and *k*_*deph*_ is identifiable and not the individual parameters. Because the individual parameters characterize processes that are much faster than the time-scales of production of our observer molecule GFP and the imaging system in the plate reader, the only information that can be passed onto the observer molecule is the net outcome of NRI protein’s phosphorylated state. Phosphorylation and dephosphorylation are processes that compete against each other to increase the amount of NRI^P^ and NRI concentration in the system, respectively. Thus, by observing the amount of NRI^P^ in the system and knowing the concentration of *NRI*^*tot*^, we can deduce the net outcome of the battle, i.e. the ratio *k*_*ph*_/*k*_*deph*_. Notice that without knowledge of *NRI*^*tot*^, we would be unable to estimate *k*_*ph*_/*k*_*deph*_. This again illustrates the importance of having additional experimental inputs for perturbing the system. Even though there is only one output molecule GFP, we are able to infer three distinct parameters that represent processes from three different time-scales: catalytic synthesis of protein, formation and disassociation of the DNA-activator complex, and phosphorylation/dephosphorylation of NRI protein.

Finally, it is worthwhile to note that the functional form of our model is consistent both quantitatively (small output residual error) and quantitatively. This suggests that our simplified model will serve as a suitable starting point for simulation studies and theoretical analysis.

## Simulations

Having all the parameters identified, we first repeated the results in Figure 2 in simulation. As shown in Figure 5A and 5B, the control circuit suffered from the retroactivity when the load was added. In contrast, the insulator circuit could attenuate the retroactivity significantly. Both the transfer curves and the fold-change comparison are consistent with the experimental results.

**Figure 5.**
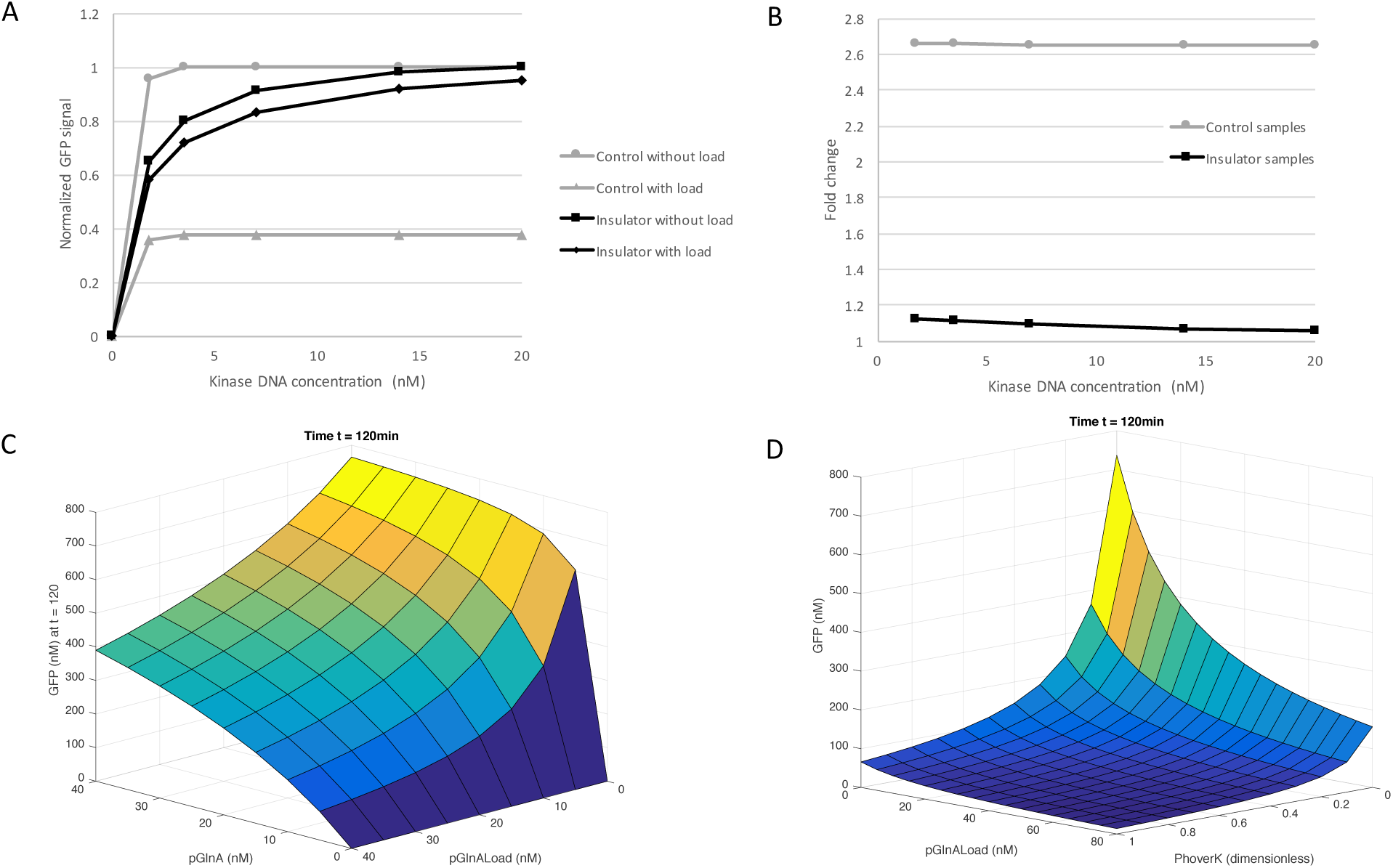
Simulation results of the PBI circuit based on the parameters identified above. **A**: Simulation results of the transfer function curves for controls and insulators with or without load DNA are consistent with the experimental results. **B**: Simulation results of the fold changes of the samples without load over the ones with load are also consistent with the experimental results. **C**: Simulation results of GFP expressions as functions of two DNA inputs. Initial condition: DNA pGlnA-GFP ranging from 0 to 40 nM and DNA pGlnA-RFP (load) ranging from 0 to 40 nM, with 68 nM protein *NRI*^*tot*^ and *Ph*^*tot*^/*K*^*tot*^ = 0. Simulation time is 120 min. D: Simulation results of GFP expressions as functions of DNA load and *Ph*^*tot*^/*K*^*tot*^. Initial condition: DNA pGlnA-RFP (load) ranging from 0 to 80 nM and *Ph*^*tot*^/*K*^*tot*^ varying between 0 and 1, with 20 nM DNA pGlnA-GFP and 68nM protein*NRI*^*tot*^. Simulation time is 120 min.

Then we wanted to know how would different initial conditions affect the output. We swept different initial conditions for the reporter pGlnA-GFP, the load pGlnA-RFP and phosphatase over kinase ratio (*Ph*_*tot*_/*K*_*tot*_) and results were shown in Figure 5C and 5D. In Figure 5C, we swept [p_GlnA_-GFP] and [p_GlnA_-RFPload]. As we can see in expression. And more load DNA brings down the GFP level, which is a result of retroactivity as load DNA competes with reporter DNA for transcription and translation resources, such as RNA polymerases and ribosomes. Load the surface plot, more reporter DNA results in more GFP DNA or mRNA sequesters those resources away from reporter DNA or mRNA, ending up with less reporter protein made. In Figure 5D, we changed the concentration of the load p_GlnA_-RFP and the ratio of Phtot/Ktot. As we mentioned above, when certain amounts of phosphatase and kinase are added, the PBI circuit will have a high gain because of phosphorylation and an equally large negative feedback by dephosphorylation. As a result, retroactivity from downstream load will be attenuated. The simulation results in Figure 5D just showed the exact same idea. By tuning the ratio, we can effectively achieve the same absolute GFP expression level with different absolute load amount. Through this mechanism, the retroactivity is largely attenuated.

## Conclusion

In this work, we investigated the structural identifiability of the phosphorylation-based insulator when implemented in a transcription-translation cell free expression system. We showed that the retroactivity exists in the TX-TL system and the PBI circuit can attenuate the retroactivity significantly. Then we considered a complex model that provided an intricate description of all chemical reactions involved in the PBI circuit. Next, leveraging specific physiologically plausible assumptions, we derived a rigorous simplified model that captures the output dynamics of the phosphorylation-based insulator. We performed standard system identification analysis and determined that the model is globally identifiable with respect to three critical parameters: the catalytic rate associated with the downstream system *k*_*cat*_, an internal parameter in the downstream system characterizing formation of the activator-DNA complex *K*_*M*_ and *k*_*ph*_/*k*_*deph*_, a ratio describing the intrinsic balance of phosphorylation and dephosphorylation in the PBI circuit. Specifically, we showed that these three parameters were identifiable only when the system was subjected to specific perturbations. We performed these experiments and estimated the parameters. Our experimental results suggest that the functional form of our simplified model is sufficient to describe reporter dynamics and enable parameter estimation. Besides, our simulations results based on the parameters estimated using above methods confirmed our conclusions from experimental data and previous theoretical predictions. These *in silico* results also showed the power of computational biology and its future applications in guiding biological experiments and synthetic biocircuits design. In general, this research illustrates the utility of the TX-TL cell free expression system as a platform for system identification, as it provides extra control inputs for parameter estimation that typically are unavailable *in vivo*. Future work will investigate the theoretical utility of the TX-TL system as a platform for system identification, parameterization of more complex systems, and the robustness and sensitivity of the phosphorylation-based insulator using our derived model.

## ASSOCIATED CONTENT

### Materials and Methods

#### Plasmids and linear DNAs

DNA and oligonucleotides primers were ordered from Integrated DNA Technologies (IDT, Coralville, Iowa). Plasmids in this study were designed in Geneious 8 (Biomatters, Ltd.) and were made using standard golden gate assembly (GGA) protocols and maintained in a KL740 strain if using an OR2- OR1 promoter (29°C) or a JM109 strain for all other constructs. Plasmids were mini prepped using Qiagen mini prep kit. BsaI-HF (R3535S) enzyme used in GGA was purchase from New England Biolabs (NEB). Linear DNAs were made by PCRing protein expression related sequences out of GGA constructs using Phusion Hot Start Flex 2X Master Mix (M0536L) from NEB. Rapid assembly procedures were based on [9].Before use in the cell-free reaction, both plasmids and PCR products underwent an additional PCR purification step using a QiaQuick column (Qiagen), which removed excess salt, and were eluted and stored in deionized water at 4°C for short-term storage and −20°C for long-term storage. All the plasmids used in the work can be found on https://www.addgene.org/.

#### TX-TL reactions and fluorescence measurements

TX-TL reaction mix was prepared and set up according to previous JOVE paper [7]. TX-TL reactions were conducted in a volume of 10 μL in a 384-well plate (Nunc) at 29°C, using a three-tube system: extract, buffer, and DNA. For deGFP, samples were read in a Synergy H1 plate reader (Biotek) using settings for excitation/emission: 485 nm/525 nm, gain 61. All samples were read in the same plate reader, and for deGFP relative fluorescence units (RFUs) were converted to nM of protein using a purified deGFP-His6 standard. Unless otherwise stated, end point measurements are after 2 h of expression at 29°C.

#### Computational models and simulations

Data analysis and fitting, model building and simulations were conducted in MATLAB (R2015b, The MathWorks, Inc.) software.

## Funding Sources

This work was supported by Office of Naval Research (ONR) Multidisciplinary University Research Initiatives (MURI) Program (Grant number: N00014-13-1-0074) and Air Force Office of Scientific Research (Grant number: FA9550-14-1-0060).

## Notes

R.M.M. has ownership in a company that commercializes the cell-free technology utilized in this paper. All the other authors claim no competing interest.

## ACKNOWLEDGMENT

We would like to thank Kayzad Soli Nilgiriwala and Domitilla Del Vecchio of MIT for providing the original plasmids and helpful suggestions. We would like to thank Murray lab members for useful discussions and suggestions.

## Supplementary Figures

**Supplementary Figure S1.**
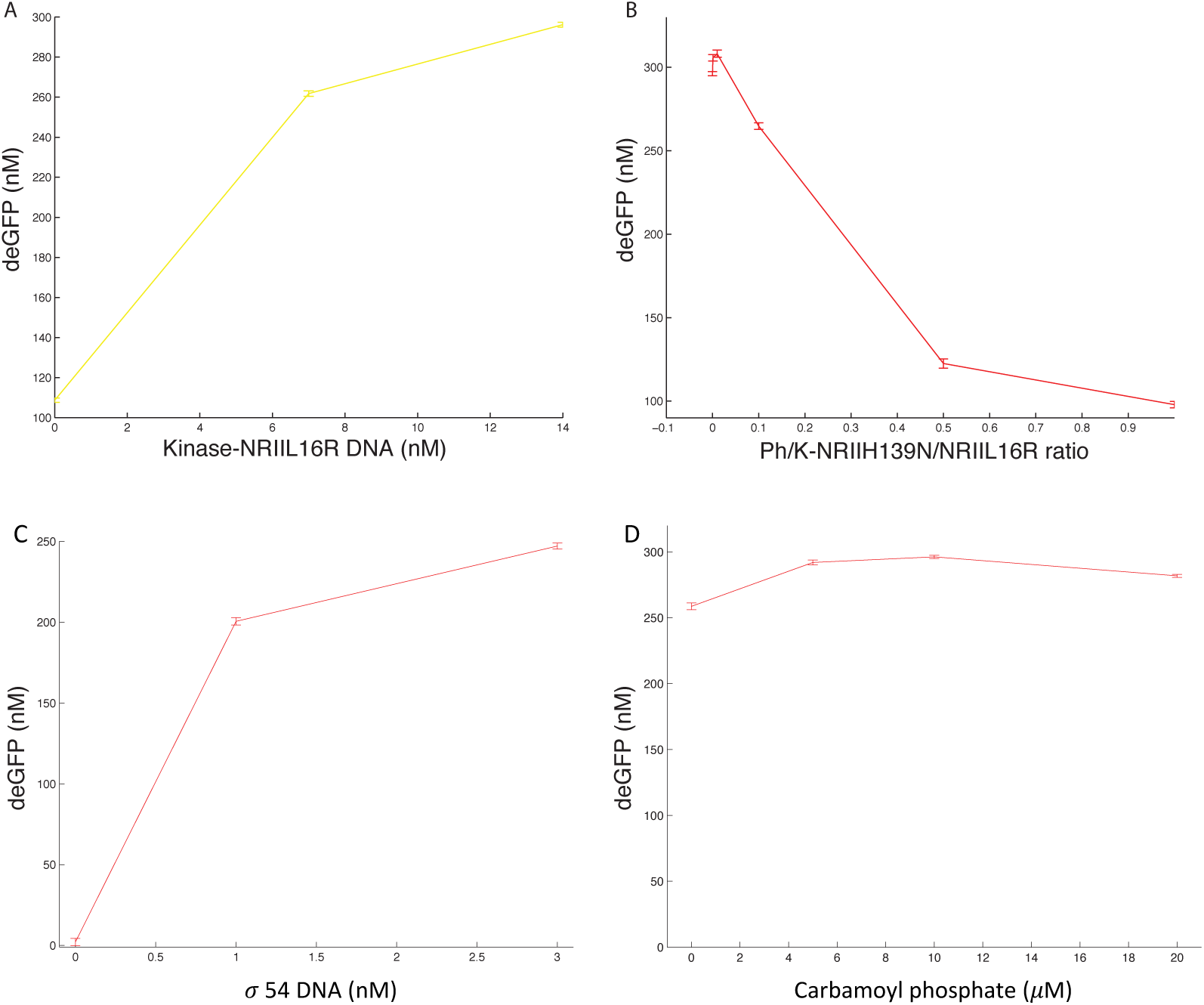
Demonstration of the phosphorylation cycle in TX-TL. Before testing the insulation capability of this circuit, we need to demonstrate that phosphorylation and dephosphorylation work in TX-TL. **A:** While keeping the phosphatase at 0 and increasing the concentrations of kinase NRIIL16R linear DNAs, we observed higher pGlnA-deGFP expression as a result of more NRI^P^. **B:** When we kept the kinase concentration constant and increased the concentrations of NRIIH139N linear DNAs (higher Ph/K ratio), less pGlnA-deGFP was expressed as a result of dephosphorylation of NRI^P^. These results suggested that the phosphorylation cycle worked in TX-TL system. **C:** We also found that σ54 DNA was required as there was little to none σ54 in the TX-TL cell extract for σ54- dependent promoter pGlnA to work. While keeping kinase, phosphatase and pGlnA-deGFP constant, adding more σ54 DNA significantly increase the expression of deGFP. **D:** We also found that additional phosphate source was recommended but not essential; as shown in the panel, increasing carbamoyl phosphate concentrations did not significantly affect the expression of deGFP. All the deGFP fluorescences have been converted to nM using purified deGFP calibration data on Bioteks.

**Supplementary Figure S2.**
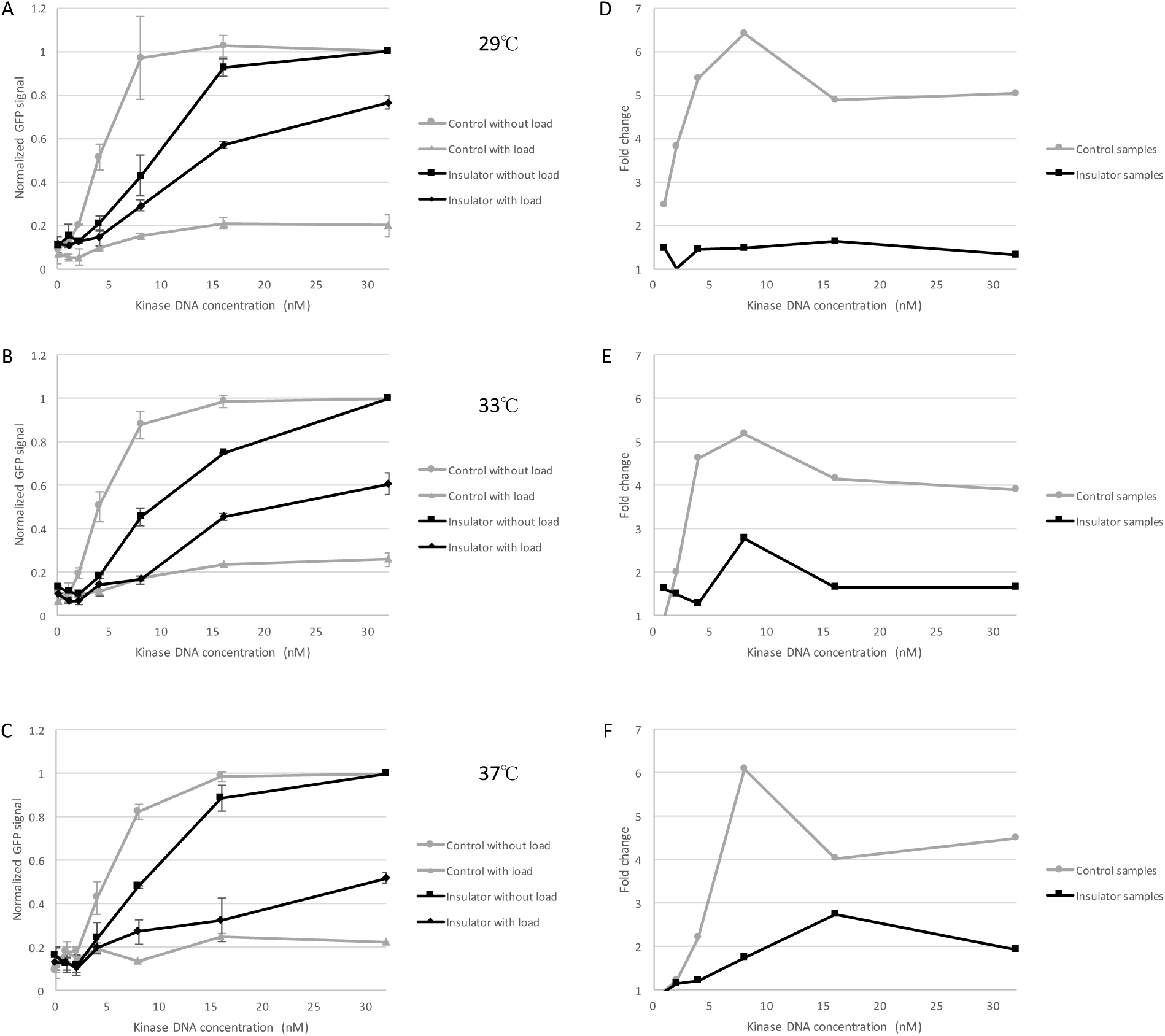
Test the temperature sensitivity of the PBI circuit at three different temperatures, 29°C, 33°C and 37°C. **A, B, C:** Transfer function curves for controls and insulators with or without load DNA at three different temperatures. **D, E, F:** Fold changes of the samples without load over the ones with load at three different temperatures.

**Supplementary Figure S3.**
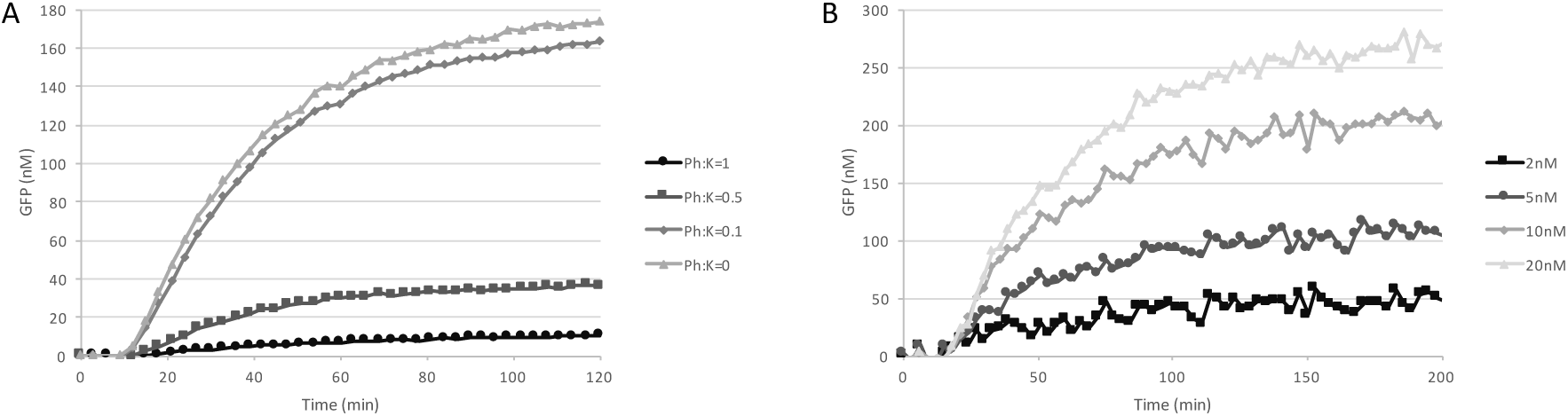
Linear regression to estimate the GFP production rate. **A:** Plot of GFP expression while varying the *Ph/K* ratio. Curves from t = 0 to *τ_max_* = 30 min were used to estimate the slope of GFP. **B:** Expression dynamics of GFP for varying amounts of pGlnA with *pLac — Ph* = 0 *nM*. These curves enable the estimation of *dGFP/dt* for *t* ≥ *τ_max_* = 30 *min*.

